# LANA-Dependent Transcription-Replication Conflicts and R-Loops at the Terminal Repeats (TR) Correlate with KSHV Episome Maintenance

**DOI:** 10.1101/2025.03.10.642343

**Authors:** Asim Asghar, Olga Vladimirova, Asher Sobotka, James Hayden, Jayamanna Wickramasinghe, Jayaraju Dheekollu, Moeko Minakuchi, Maureen E. Murphy, Kazuko Nishikura, Paul M. Lieberman

**Affiliations:** The Wistar Institute, Philadelphia, PA 19104

**Keywords:** R-loops, H3pS10, episome maintenance, LANA, KSHV, DNA virus

## Abstract

Transcription-replication conflicts frequently occur at repetitive DNA elements involved in genome maintenance functions. The KSHV terminal repeats (TR) function as the viral episome maintenance element when bound by the viral encoded nuclear antigen LANA. Here, we show that transcription-replication conflicts occur at or near LANA binding sites in the TR. We show by proximity ligation assay (PLA) that PCNA and RNAPII colocalize with LANA-nuclear bodies (LANA-NBs). Using DNA-RNA-IP (DRIP) assays with S9.6 antibody, we demonstrate that R-loops form at the TR. We find that these R-loops are also associated with histone H3pS10 a marker for R-loops associated with transcription-replication conflicts. Inhibitors of RNA polymerase eliminated LANA binding to the TR, along with the loss of R-loops and activation associated histone modifications, and the accumulation of heterochromatic marks. We show that LANA can induce all of these features on a plasmid containing 8, but not 2 copies of the TR, correlating strongly with episome maintenance function. Taken together, our study indicates that LANA induces histone modifications associated with RNA and DNA polymerase activity and the formation of R-loops that correlate with episome maintenance function. These findings provide new insights into mechanisms of KSHV episome maintenance during latency and more generally for genome maintenance of repetitive DNA.

**Importance:** KSHV latent infection is responsible for Kaposi’s Sarcoma (KS) and Pleural Effusion Lymphoma (PEL). KSHV latency and persistence depends on LANA binding to the terminal repeats (TR). We show that LANA binding promotes the formation of R-loops associated with transcription-replication conflicts and histone H3pS10 at the KSHV terminal repeats. These epigenetic features depend on active RNA polymerase at the TR and correlate strongly with KSHV episome maintenance function. The findings suggest a novel mechanism of chromatin structural maintenance dependent on LANA binding at the TR during KSHV latency.

## Introduction

Genome maintenance elements, such as telomeres and centromeres, typically consist of arrays of repetitive genetic elements recognized by sequence-specific factors that assemble into a higher-order structure and confer functions essential for genome protection, replication and segregation during cellular division [1, 2]. The terminal repeats (TRs) of the Kaposi Sarcoma-Associated Herpesvirus (KSHV), also known as HHV8, consists of a highly GC-rich ∼800 bp repeat element that serves as a selective binding site for the viral protein LANA and function as both origin of DNA replication and episome maintenance element in latently infected host cells [3–5]. Like human telomeres and centromeres, the KSHV TR provides genome protection and higher-order structure that can influence transcription both locally and at distal genetic sites [6].

KSHV is an oncogenic human gammaherpesvirus that is responsible for Kaposi’s sarcoma (KS) and pleural effusion lymphomas associated with HIV AIDS. KSHV is also a causative agent of multicentric Castleman’s Disease (MCD) [7, 8]. KSHV establishes a long-term latent infection in B-lymphocytes where it persists as a multi-copy covalently closed minichromosome of ∼170,000 kb, referred to as the viral episome [9, 10]. Cellular chromatin assembles on the KSHV episomes to repress most of the viral genes and maintain a stable latent state [11, 12]. Maintenance of the latent state requires the expression of the viral-encoded Latency Associated Nuclear Antigen (LANA). LANA regulates the latency program of the virus by binding to three sites within the 800 bp TR unit that are repeated up to 30 copies per genome [13–15]. Episome maintenance function requires at least 8 copies of the TR, while DNA replication requires only the 2 LANA binding sites within each single TR unit [9]. LANA also forms large foci at the TR of each viral episome suggesting that a higher-order structure is an important regulatory feature of the KSHV episome maintenance function [9, 16, 17].

LANA consists of two major subdomains important for its functional activity in viral episome maintenance. The C-terminal domain of LANA comprises the sequence-specific DNA binding domain (DBD) which can form higher order oligomeric structures on DNA [14, 18]. The N-terminal domain enables tethering to the metaphase chromosome [19, 20] and can interact directly with an acidic patch in histones H2A/B which correlates strongly with viral replication and episome maintenance activity [21, 22]. The N-terminal domain can also interact with components of the transcriptional co-activator complex MLL1, which can modify both histones and RNA polymerase II to promote active transcription [23]. Cellular replication factors, including origin binding proteins (ORCs) and minichromosome maintenance (MCMs) responsible for the initiation of DNA replication are found to be enriched at LBS sites in the TR [11] and LANA can co-immunoprecipitate with proliferating cell nuclear antigen (PCNA)[24]. Nucleosomes are also found to assemble around the LBS in the TR, and these are marked by specific modifications that vary across the cell cycle [11, 25, 26]. Whilst the bulk of the KSHV genome is marked by repressive H3K27me3 and H3K9me3, the TR is marked by H3K4me3 and H3K27ac, characteristic of an open chromatin structure and enhancer function. Indeed, it has been shown the TR has enhancer and transcriptional activity [6, 25]. LANA can also interact with other chromatin regulatory factors, including BRD4, ADNP, and CHD4 [6, 25]. The histone H3.3 chaperone DAXX has also been found to bind LANA and colocalize with LANA-nuclear bodies on KSHV episomes [17, 27]. Although not directly interacting with LANA, the chromatin organizing factors CTCF and cohesin are enriched near the LANA binding sites (LBS) within the TR [28, 29]. Precisely how these factors work together to orchestrate the LANA-dependent episome maintenance function at the KSHV TR remains only partly understood.

Replication and transcription machinery may translocate along the same DNA template, often in opposing directions and at different rates leading to transcription-replication conflicts (TRCs) [30, 31].. High GC content and repetitive DNA can further lead to challenges for both transcription and DNA replication that further promote TRCs. Tightly bound proteins, such as arrays of LANA binding sites in the TR may also promote replication fork pausing that can also lead to TRCs. Unresolved TRCs can lead to catastrophic genome instability and cell cycle arrest. On the other hand, some TRCs may provide specialized features such as formation of higher-ordered chromatin structures that facilitate the maintenance of repetitive genetic elements. Here, we show that LANA binding to the KSHV TR leads to the colocalization of transcription and replication machinery. This leads to the formation of an RNA-DNA hybrid, or R-loop, that is associated with histone H3pS10, a signal typically observed at transcription-replication conflicts and also on condensed mitotic chromosomes. We propose that LANA induces these epigenetic features at the KSHV TR to facilitate both dynamic activity and protective condensation necessary for stable maintenance of the viral episome during latency.

## Results

### Colocalization of transcription and replication factors at the KSHV TR

Previous studies have identified histone modifications and epigenetic factors that localize with LANA at the KSHV TRs. We used ChIP-qPCR to confirm that H3K27ac, H3K4me3, CTCF, and RAD21 are enriched at the LBS region of the TR in latently infected BCBL1 and iSLK cell lines (**Supplementary Fig. 1**). We next asked whether DNA replication factors associated with active replication forks, including MCMs and PCNA colocalize with RNA polymerase phospho-isoforms associated with either elongating (pS2) or promoter proximal pausing (pS5) are also colocalized at the TR (**Fig. 1**). ChIP-qPCR demonstrated that PCNA, MCM2, along with RNA Pol II pS2 and pS5 are all selectively enriched at TR relative to control IgG and relative to other regions of the KSHV genome, such as ORF45 and ORF75 (**Fig. 1A-C).** To determine if replication and transcription factors were spatially and temporally colocalized with LANA at the TR, we performed proximity ligation assays (PLA) combined with immuno-fluorescence imaging in KSHV infected iSLK cells (**Fig. 1D** and **E**). We performed PLA with antibodies to PCNA as a marker for DNA replication forks and with either RNAPII pS2 or pS5. Colocalizations detected by PLA were visualized by green fluorescence and were only observed when both pairs of antibodies were present, but not in control samples lacking primary antibodies (**Fig. 1C**). Numerous colocalizations were observed for PCNA with both pS2 and pS5, and are likely to represent many expected sites of co-incident transcription and replication throughout the cellular genome. PLA signals were then assayed for their colocalization with RFP-LANA nuclear bodies in iSLK cells latently infected with KSHV genomes expressing RFP-LANA. We found that ∼30% of LANA-NBs colocalized with RNAPII pS5 and ∼15% colocalized with RNAPII pS2. This suggests that nearly one third of LANA-NBs have coincidence of replication forks (PCNA) with RNA polymerase II pS5 and to a lesser extent pS2.

**Fig. 1.**
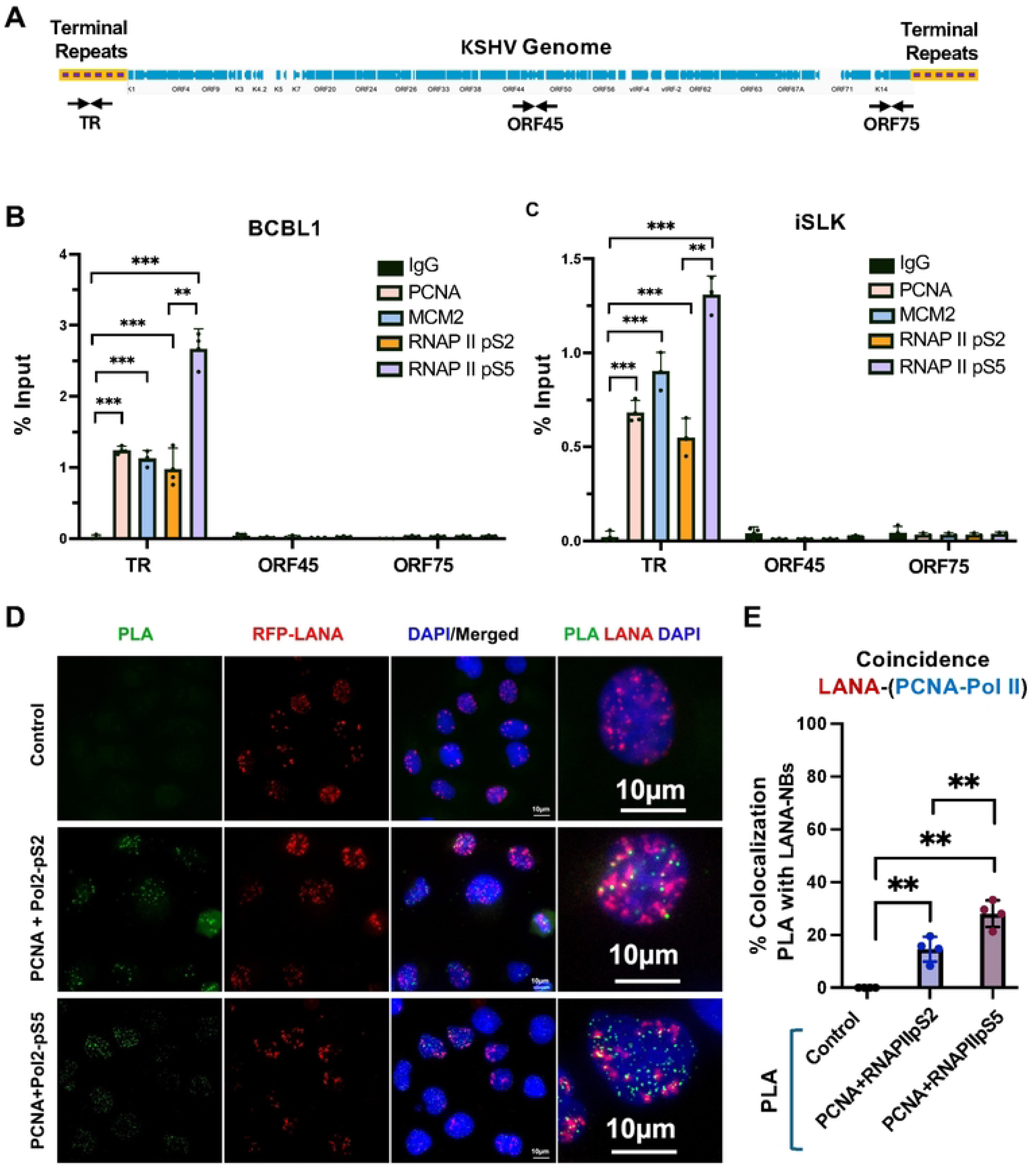
Colocalization of transcription and replication machinery with LANA at KSHV TR. **A**. Schematic of the KSHV genome showing the terminal repeats (TR) relative to the unique region open reading frames (blue) and primer positions for TR, ORF45, and ORF75. **B and C**. ChIP-qPCR for RNAP II pS5, RNAPII pS2, MCM2, PCNA or control IgG assayed at the TR, ORF45 or ORF75 loci in BCBL1 (B) or iSLK (C) cell lines. ** p<.01, *** p<.001, student 2-tailed t-test, n=3 biological replicates. **D**. Representative IF microscopy image showing PLA (green) for PCNA+RNAP II-pS2 or PCNA+RNAPII-pS5, RFP-LANA (red), Dapi (blue) and merge. 60X. **E.** Quantification of IF images showing % colocalization of PLA signal with LANA-NBs. ** p<.01, pairwise Anova.

### R-Loops form at TR and colocalize with LANA-NBs

Transcription-replication conflicts frequently coincide with R-loops formed by unresolved RNA-DNA hybrids [32, 33]. R-loops have been previously identified in the KSHV TR as well as the ORF16 locus [34]. We confirmed these findings using the S9.6 monoclonal antibody for DNA-RNA IP (DRIP) assay. DRIP revealed that R-loops were detectable at the KSHV TR, but not at other viral loci for ORF45 and ORF75 for BCBL1 (**Fig. 2A**) and BC-1 (**Supplemental Figure 2**). To verify that S9.6 was recognizing RNA-DNA hybrids, we treated the immunopurified material with RNAse H, which eliminated the signal from the TR region, as well as from the positive control ORF16 region (**Fig. 2B**). To determine if R-loops colocalize with LANA at LANA-NBs, we performed PLA with antibodies to LANA and S9.6 (**Fig. 2C**). Nuclear dots were detected only when both antibodies were present, indicating that LANA is in close proximity to S9.6 detected R-loop structures.

**Fig. 2.**
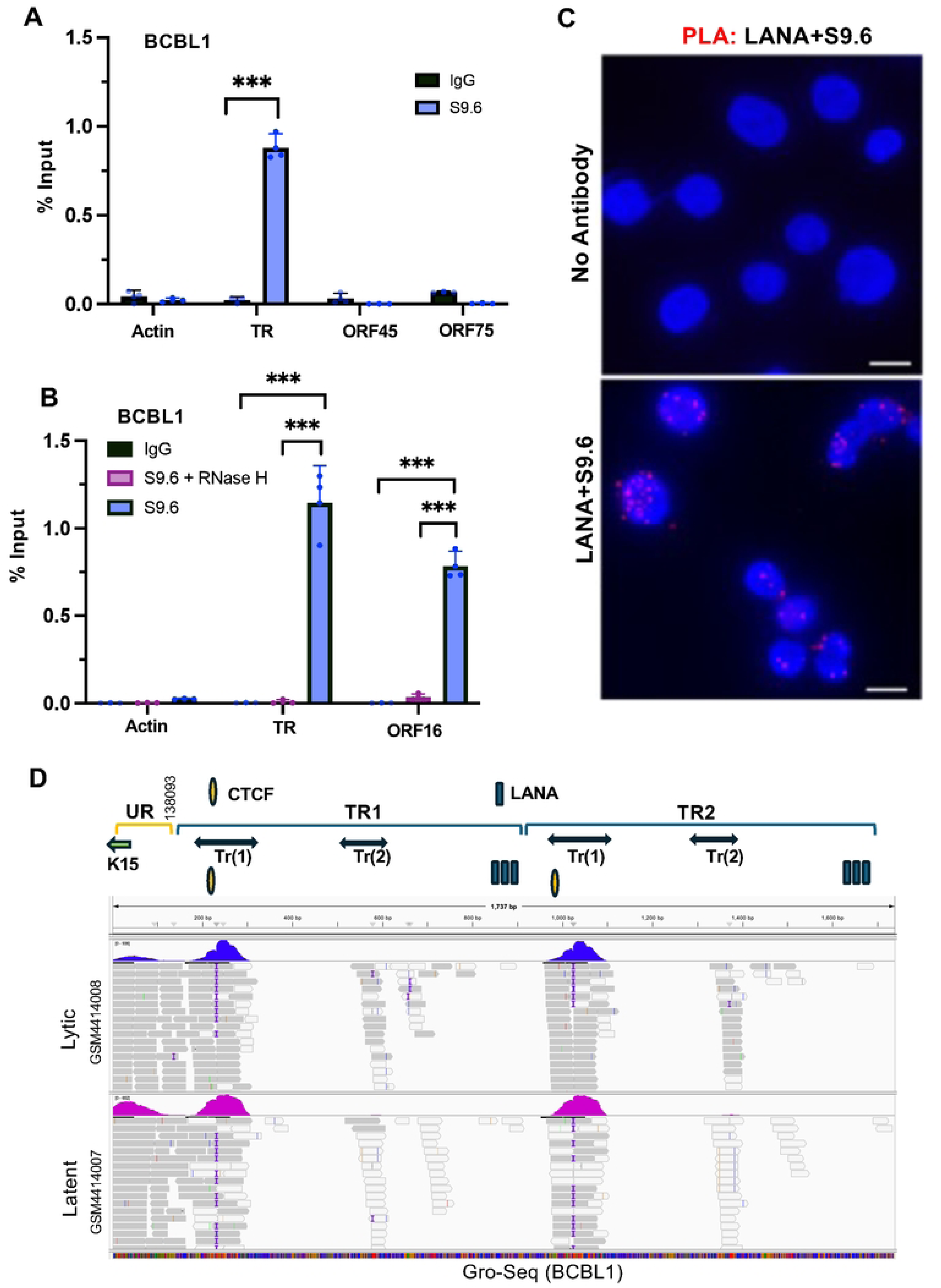
R-loop formation at the KSHV TR. **A.** DRIP assay with BCBL1 cells using S9.6 (blue) or control IgG (black) assayed with primers for cellular actin, KSHV TR, ORF45 or ORF75. **B.** Same as in panel A, except for KSHV TR or ORF16 and with specificity control RNase H treatment (pink). *** P<.001, student two-tailed t-test. **C.** PLA analysis of LANA+S9.6 (red signal) in BCBL1 cells. Dapi (Blue). No antibody control is shown in top panel. **D.** IGV screen shot of RNA transcripts mapped to KSHV TR region using public data sets for GROseq in BCBL1 in latent or lytic conditions (GSM4414007, GSM4414008). The reference map consists of a small region of the unique region with K15 and 2 copies of the TR. TR transcripts Tr(1) and Tr(2) are indicated above.

To determine if R-loop RNA mapping to the TR could be identified, we analyzed a public Gro-Seq [35] and total RNA-seq [36] datasets for BCBL1. The SRR data set for GRO-Seq (**Fig. 2D**) and total RNA-seq (**Supplemental Figure 2B**) both detect a series of transcripts that mapped to the KSHV TR. We identified two transcripts, referred to Tr(1) and Tr(2) that mapped to regions within the TR. The more prominent transcript Tr(1) appears to initiate near LANA binding sites and terminate near the adjacent CTCF binding site. These findings indicate that small RNA fragments are generated within the TR.

### H3pS10 localizes to TR and a subset of LANA-NBs

R-loops linked to transcription-replication conflict have been reported to generate phosphorylated histone H3 (H3pS10) during interphase [37]. To determine whether the R-loops at TR were associated with histone H3pS10, we assayed the localization of H3pS10 with TR by ChIP-qPCR (**Fig. 3A**). We found that H3pS10 was highly enriched at TR, but not detectable at ORF45 or ORF75. To assess the cell cycle dependence of these colocalizations we fractionated cells according to their stage of the cell cycle using centrifugal elutriation and then assayed these by ChIP-qPCR (**Fig. 3B**). We found that H3pS10, along with MCM and RNAPII pS2 were highly enriched at the TR in G1 and G1/S. MCM and H3pS10 were observed also in G2 and G2/M. At the control region for ORF16, which also forms an R-loop, we found only RNAPII enrichment in G1 and G1/S phase of the cell cycle. (**Fig. 3C**). This suggests that H3pS10 associates with the R-loop at the TR during G1 and G2/M, but does not associate with the ORF16 R-loop at any stage of the cell cycle.

**Fig. 3.**
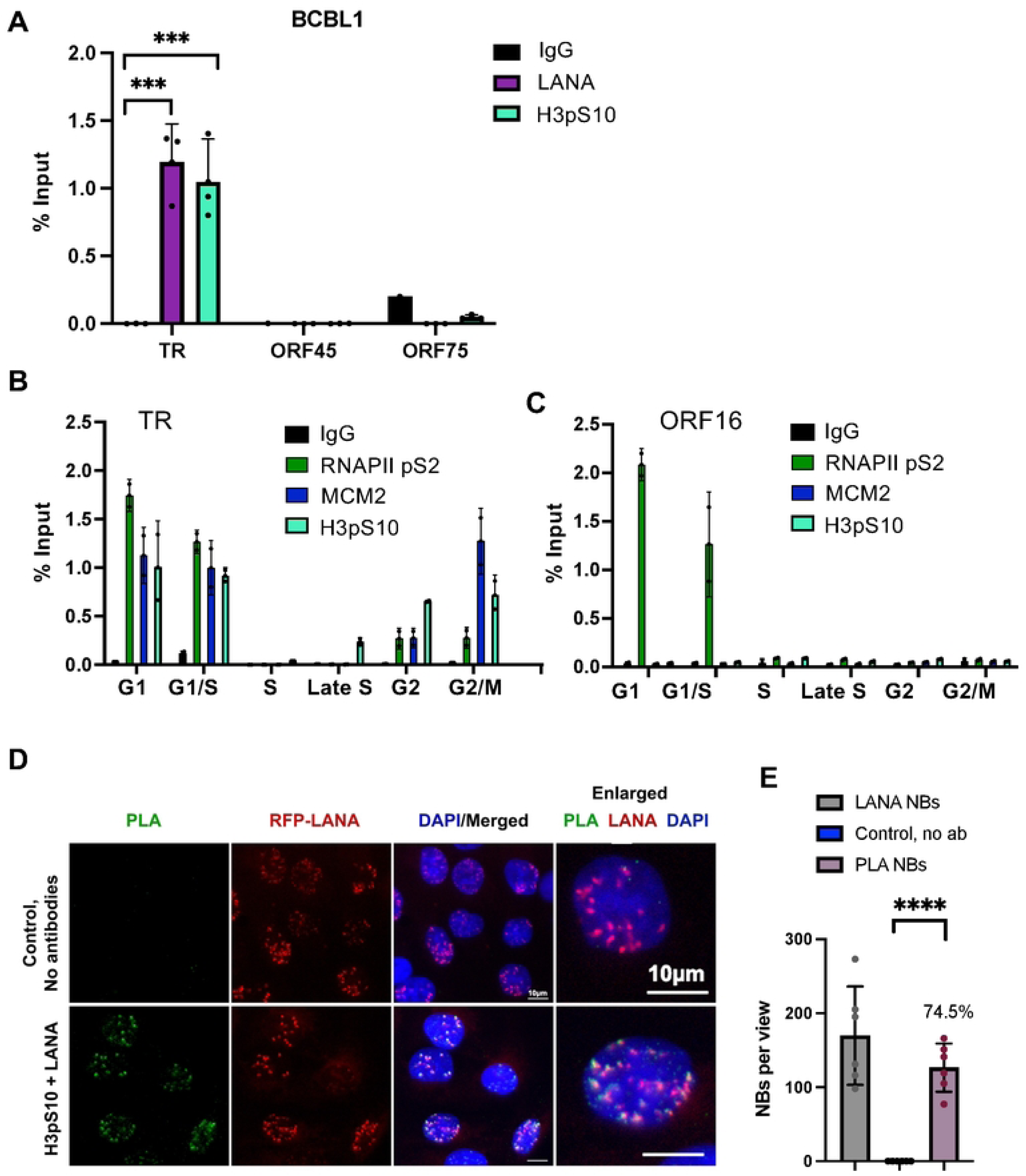
Cell cycle dependent accumulation of H3pS10 at the TR. **A**. ChIP-qPCR for LANA, H3pS10 and IgG at the TR, ORF45, ORF50, and ORF75 loci of KSHV in BCBL1 cells. **B-C**. ChIP-qPCR for RNAPII pS2, MCM, H3pS10 during different stages of the cell cycle using centrifugal elutriation for G1, G1/S, S, Late S, G2, and G2/M. ChIP-qPCR was analyzed at the TR (panel B) or at the ORF16 locus (panel C). **D.** IF of single cell showing H3pS10 (red), LANA (green), Dapi (blue) and merge. **E**. PLA for H3pS10+LANA (green), with RFP-LANA (red), Dapi (blue) and merge in RFP-LANA iSLK cells. No-antibody control shown in top panels. **F.** Quantification of data for representative IF images shown in panel E. **** p<.00001, **p <0.01 using two-tailed t test with Mann-Whitney test and Welch’s correction.

To further validate the association of H3pS10 with TR, we performed PLA with H3pS10 and LANA (**Fig. 3D**). These PLA signals were found to colocalize with a significant fraction (∼74.5%) of LANA-NBs and were strictly dependent on primary antibody (**Fig. 3E**).

Colocalization of H3pS10 with LANA-NBs and replication centers were further analyzed by IF with H3pS10 and PCNA (**Fig. 4** and **Supplemental Figures 3-5**). We found that PCNA colocalized with ∼29.7% of LANA-NBs, H3pS10 colocalized with 26.9% of LANA-NBs, and H3pS10 colocalized with 52.9% of PCNA foci. All three components are colocalized at ∼21.7% of LANA-NBs in iSLK cells (**Figure 4G** and **H, and Supplemental Figures 3** and **5**). Higher resolution confocal microscopy was used to further characterize features of LANA-NBs that contain both H3pS10 and PCNA (**Supplemental Figures 4, 5 and Movie 1**). LANA-NBs with coincident H3pS10 tended to be larger substructures compared to LANA-NBs lacking these colocalizations. These images indicate that a large subnuclear domain encompassing LANA is frequently colocalized with both PCNA and H3pS10.

**Fig. 4.**
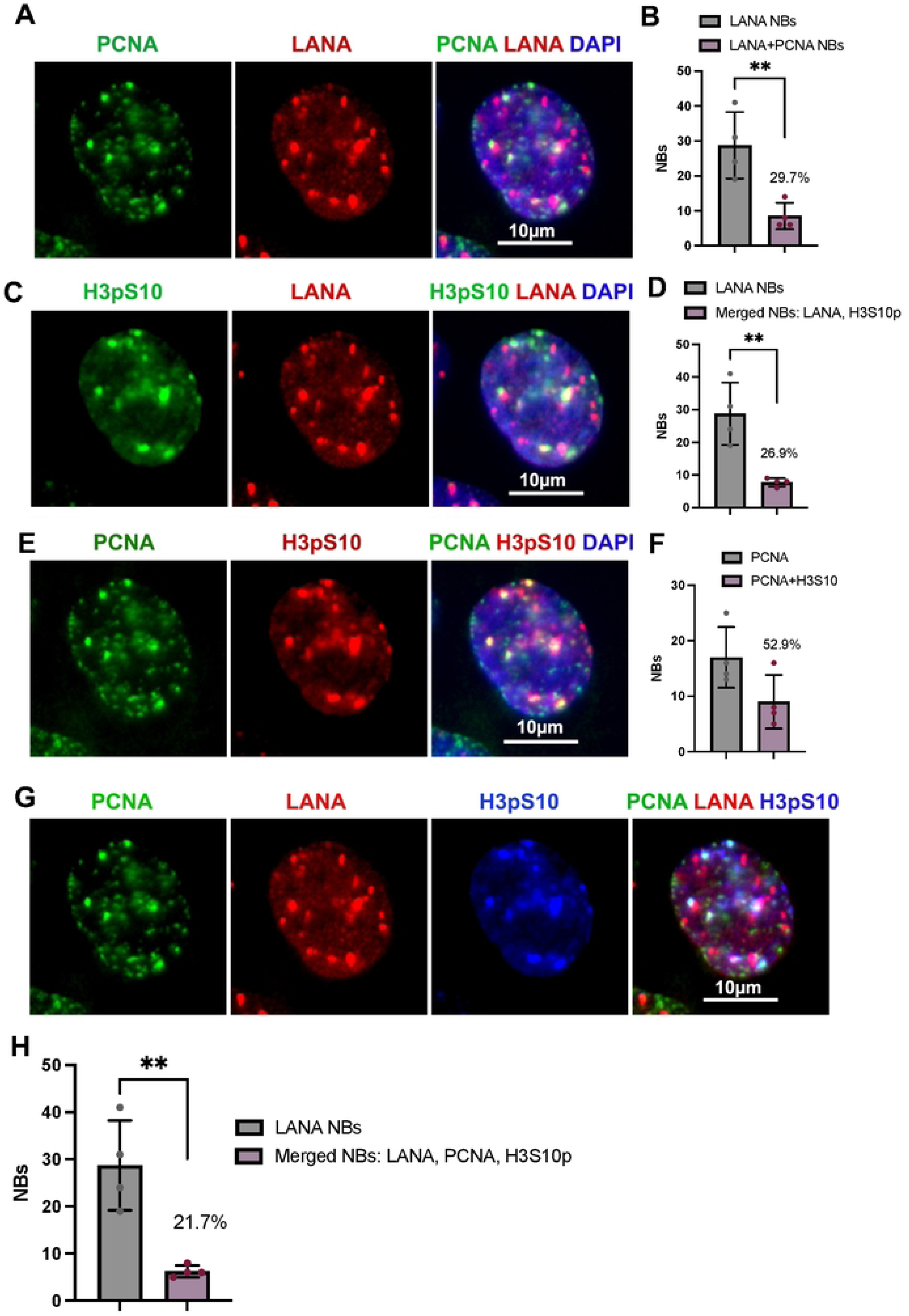
IF colocalization of PCNA-H3pS10-LANA in iSLK cells. **A**. PCNA (green), LANA (red), Dapi (blue). **B**. Quantification of the percent of LANA-NBs colocalized with PCNA. **C.** H3pS10 (green), LANA (red), Dapi (blue). **D.** Quantification of the percentage of LANA-NBs colocalized with H3pS10. **E.** PCNA (green), H3pS10 (red), Dapi (blue). **F.** Quantification of the percentage of PCNA foci colocalized with H3pS10. **G**. Combined IF with PCNA (green), LANA (red), H3pS10 (blue). **H.** Quantification of the percentage of LANA-NBs colocalized with H3pS10 and PCNA. N=4, total of 35 cells, **p<0.01, student two-tailed t-test.

**Fig. 5.**
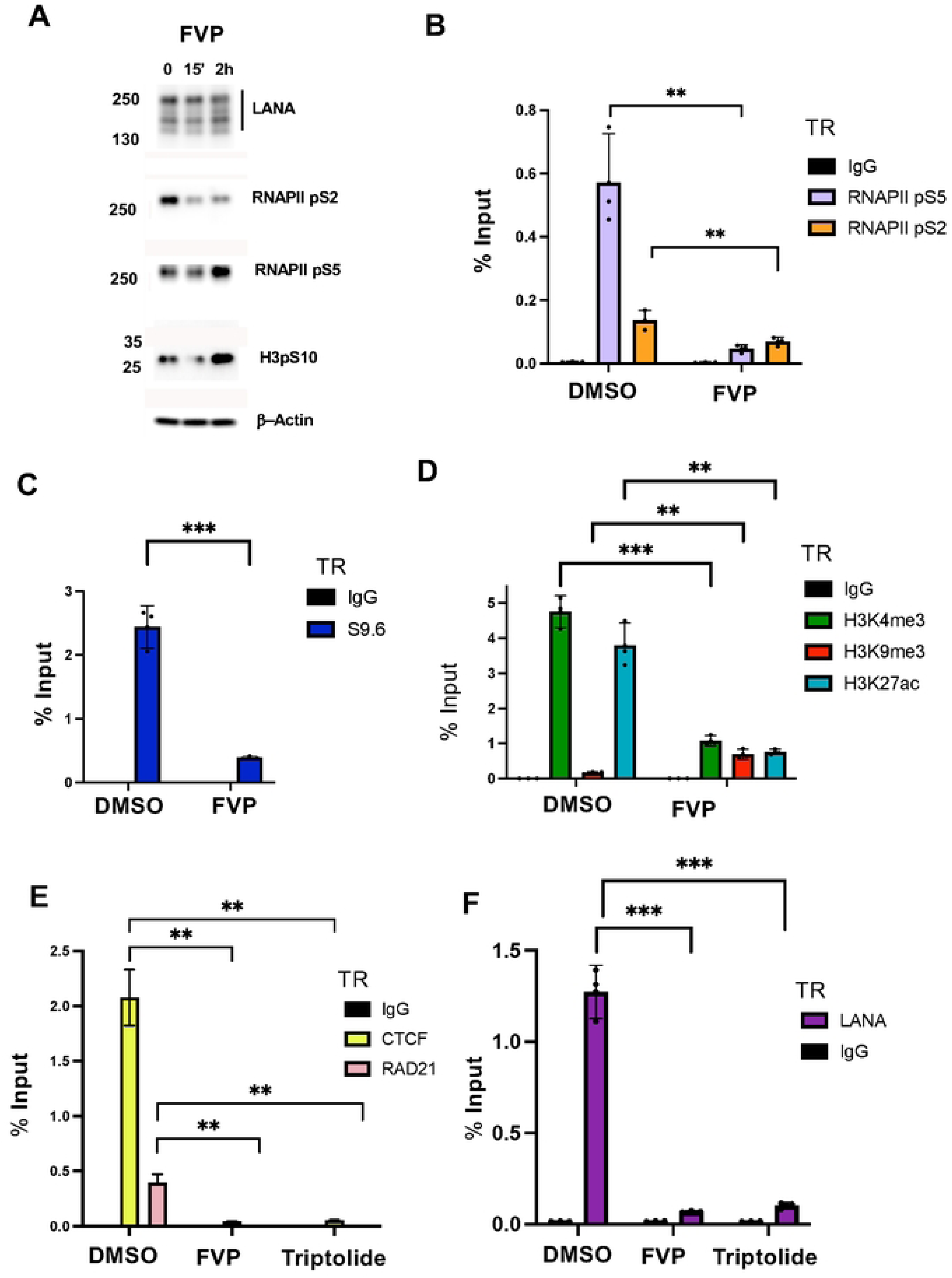
RNA polymerase inhibitors block LANA binding, R-loop formation, and epigenetic programming at the TR. **A.** Western blot of LANA, Pol2-pS2, Pol2-pS5, and b-Actin in BCBL1 cells treated with flavopiridol (FVP) for 0, 15 min or 2 hrs. **B**. ChIP-qPCR for IgG, RNAPII pS2 or pS5 at the TR in BCBL1 cells treated with DMSO or FVP for 15’. **C.** DRIP assay with S9.6 or IgG for BCLB1 cells treated with DMSO or FVP for 15’. **D.** ChIP-qPCR for H3K4me3, H3K9me3, H3K27ac or IgG in BCBL1 cells treated with DMSO or FVP for 15’. **E**. ChIP-qPCR for CTCF, RAD21, or IgG at TR in BCLB1 cells treated with DMSO, FVP, or Triptolide for 15’. **F.** ChIP-qPCR for LANA or IgG at TR in BCLB1 cells treated with DMSO or FVP for 15’ or Triptolide for 15’. ** p<.01, ***p<.001, two tailed student t-test.

### RNA polymerase II inhibitors disrupt active histones, R-loops, CTCF-cohesin and LANA binding to TR

The CDK9 inhibitor flavopiridol (FVP) rapidly inhibits RNA polymerase activity by blocking phosphorylation at S2 and preventing elongating RNA synthesis. Treatment of BCBL1 cells with FVP for 15’ or 2 hrs had minimal effects on viability, but did show loss of RNAPII pS2 by Western blot (**Fig. 5A and Supplemental Figure S6**). LANA expression pattern did not change significantly at this early time after treatment (**Fig. 5A**). As expected, RNAP II pS2 binding was significantly reduced at the TR after 15’ treatment of FVP treatment. We also found a significant loss in RNAP II pS5 (**Fig. 5B**). We next assayed the effects of FVP on R-loop formation at the TR using DRIP assay with S9.6 antibody (**Fig. 5C**). We found that 15 min of FVP resulted in a ∼6-fold reduction in DRIP signal. Thus, R-loops are also dependent on RNAP II phosphorylation and continued activity in the TR. We next assayed histone modifications associated with the TR. We found that FVP reduced active histone marks for H3K27ac and H3K4me3, but increased the signal for the heterochromatic mark H3K9me3 (**Fig. 5D**). We also found that FVP treatment reduced CTCF and RAD21 binding (**Fig. 5E**). Surprisingly, we also found that FVP treatment for 15 min led to a dramatic loss of LANA binding to the TR (**Fig. 5F**). To verify this result, we tested a second inhibitor of RNA polymerase, triptolide, that works through a different mechanism involving the degradation of RNA polymerase protein (**Supplemental Fig. S6**) [38]. Similar to FVP, triptolide treatment for 15 min also led to a drastic reduction in LANA, as well as CTCF and RAD21 binding to the TR (**Fig. 5E** and **F**). These findings indicate that inhibitors of RNA polymerase disrupt LANA binding along with R-loops and epigenetic features associated with the TR.

### Epigenetic Dependencies on LANA and TR repeat number

The functional episome maintenance activity of the TR depends on the number of terminal repeats such that 8xTR is functional but 2xTR is non-functional [39]. We therefore tested whether some of these epigenetic features depend on LANA binding and correlate with 8 but not 2 TR repeats **(Fig. 6A**). We first noted that the 8xTR was enriched for the heterochromatic H3K9me3 in the absence of LANA (**Fig. 6B**, F-Vector). Cotransfection of Flag-LANA resulted in LANA binding, along with enrichment of H3K4me3 and H3K27ac, and decrease in H3K9me3 (**Fig. 6B**, F-LANA). LANA expression also stimulated the assembly of RNAPII pS2 and pS5 (**Fig. 6C**). In contrast to the 8xTR template, the 2xTR template failed to assemble H3K27ac or H3K4me3 despite substantial binding of LANA (**Fig. 6D**). Similarly, RNAPII pS2 and pS5 failed to assemble on 2xTR while they efficiently assemble on 8xTR in the presence of LANA (**Fig. 6E**).

**Fig. 6.**
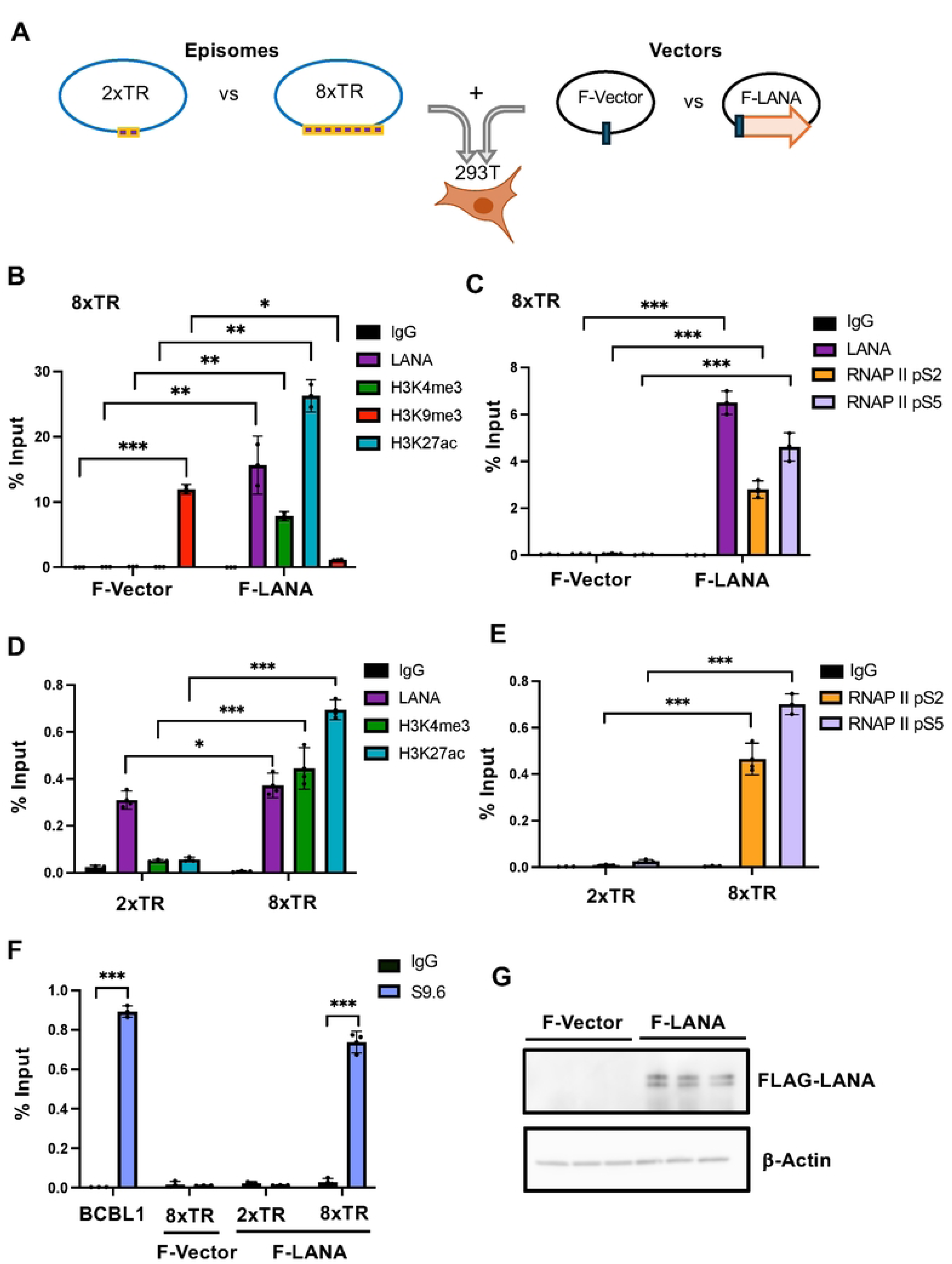
LANA and TR repeat number dependence of RNA polymerase II activation, epigenetic programming and R-loop formation at the KSHV TR. **A.** Schematic of the experimental design for co-transfection of either 2xTR or 8xTR episome templates in combination with either empty FLAG-vector (F-Vector) or vector expressing FLAG-LANA (F-LANA) into 293T cells. **B**. ChIP-qPCR of 8xTR episome template with F-Vector or F-LANA assayed for LANA, H3K4me3, H3K9me3, H3K29ac or IgG. **C**. ChIP-qPCR for 8xTR episome template with F-Vector of F-LANA assayed for LANA, RNAP II pS2, pS5, or IgG. **D**. ChIP-qPCR with 2xTR or 8xTR template in the presence of F-LANA assayed for LANA, H3K4me3, H3K27ac, or IgG. **E**. ChIP-qPCR for 2xTR or 8xTR in present of F-LANA assayed for RNAP II pS2, pS5 or IgG. **F**. DRIP with S9.6 or IgG assayed at the TR for BCBL1, 8xTR with F-Vector, 2xTR with F-LANA or 8xTR with F-LANA. **G**. Western blot of 293T cells transfected with F-Vector of F-LANA probed for FLAG or b-actin. *<.05, ** p<.01, p<.001, two tailed student t-test.

R-loops assayed by DRIP assay were also found to be dependent on both LANA and 8xTR, as they did not form on 2xTR with LANA or 8xTR lacking LANA (**Fig. 6F**). These findings indicate that R-loops, RNAPII and active histone modifications assembly at TR in the presence of LANA and correlates with functional episome maintenance activity.

## DISCUSSION

The episome maintenance of the latent KSHV genome requires coordination of DNA replication, RNA polymerase II transcription and epigenetic programming that are essential for viral genome stability and persistence. How these potentially conflicting activities are choreographed remains poorly understood. Here, we provide new evidence that during latent infection RNA polymerase and DNA replication machinery colocalize within the TR in association with RNA-DNA hybrid (R-loop) and histone modifications associated with transcription-replication conflicts (TRCs). We show that active RNA polymerase is required for many of these epigenetic features to form at the TR, including LANA and CTCF chromatin binding. Finally, we show that R-loops and active histone modifications correlate with the functional episome maintenance activity of LANA binding and multiple copies of the TR, and the prevention of default H3K9me3 heterochromatinization (**Fig. 7** model).

**Fig. 7.**
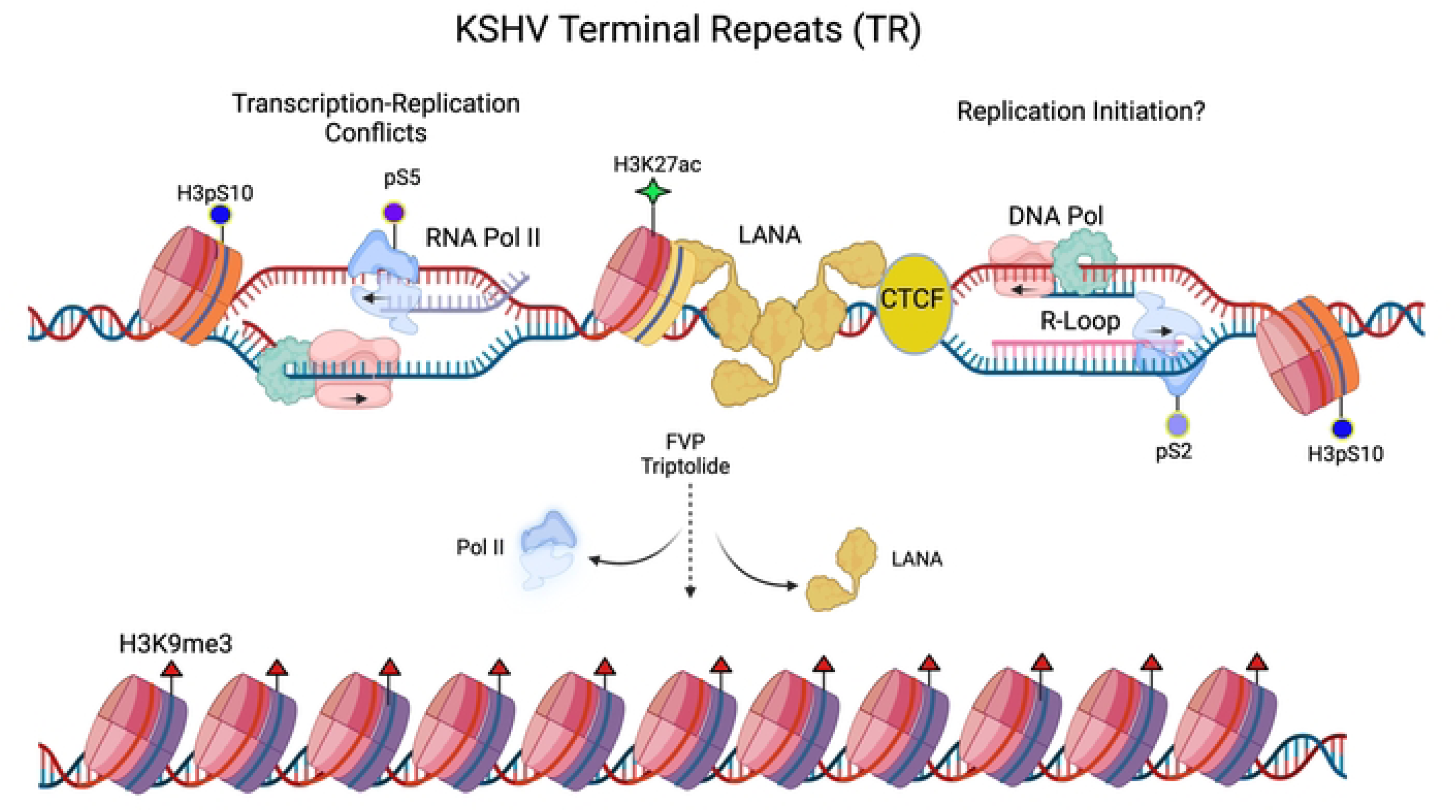
Model of LANA-dependent R-loops formed by collisions between elongating RNA polymerase II and DNA replication machinery at the TR. Inhibition of RNA polymerase leads to a loss of LANA binding, R-loops and the formation of heterochromatic H3K9me3.

### TRCs at the KSHV TR

Transcription-replication conflicts are known to be a source of R-loops that can lead to replication fork collapse and genetic instability [30]. LANA is known to recruit components of the replication machinery, including ORC, MCMs, and PCNA to the viral replication origin within the TR [11, 24]. Genetic analysis of the minimal viral replication origin identified two central pairs of LANA binding sites in the TR [15], while structural studies indicate that an additional LANA binding site is also present [14]. LANA is also known to regulate transcription across the viral and host genome [40–42], although the precise mechanism is not fully established. LANA can interact with components of the MLL1 complex that can regulate histone H3K4me3 methylation and RNA polymerase initiation through interactions with PTEFb complex [43, 44]. We found that both replication (PCNA and MCM2) and transcription factors (RNA polymerase II) associate simultaneously with LANA at the TR using PLA and cell-cycle dependent ChIP-qPCR. These findings strongly suggest that transcription and replication coincide at the TR.

### R-Loops at the KSHV TR

R-loops may form at different genetic elements with different functional or pathological consequences. R-loops can form as a result of RNA polymerase promoter proximal pausing [45] or stalling due to steric interference, such as at CTCF binding sites [46, 47]. Promoter proximal pausing can generate small R-loops due to the inhibition of CDK9-pTEFB conversion of RNAPII from pS5 to pS2. R-loops can also form when RNAPII pauses due to collisions with DNA polymerase or due to complex DNA structures, such as repetitive DNA and G-quadruplexes, that hinder polymerase processivity [48, 49]. At the KSHV TR, we found that R-loops depended on active RNA polymerase as they were inhibited by FVP (Fig. 5). R-loops were also dependent on LANA binding to the 8xTR template (Fig. 6). RNAseq analyses suggests that transcripts mapping to the TR may start and end near CTCF and LANA binding sites. Both CTCF and LANA have been implicated in interactions with either RNA polymerase II directly, or with associated initiating cofactors, such as LANA interaction with MLL1 complex [23]. ADNP, a factor that has been shown to bind LANA and associate with TR [50], is also implicated in the localization and regulation of R-loops [51]. These findings suggest that both CTCF and LANA regulate the generation of short TR-associated RNAs that remain associated with the TR as R-loops.

### H3pS10 colocalization with LANA and R-loops at the TR

H3pS10 has been found to localize to R-loops in multiple organisms, including yeast and human [37, 52]. The association of H3pS10 with R-loops was linked to replication fork stalling and chromosome condensation [37]. H3pS10 is typically associated with condensed mitotic chromosomes, but those observed with LANA-NBs and TR were found in G1 and G2 phases of the cell cycle, and also colocalized with active chromatin marks for H3K27ac and H3K4me3. H3S10 phosphorylation can also serve as a phospho-switch to activate transcription and prevent heterochromatinization at some regulatory regions, including HSV during latent infection [53, 54]. Others have found that mitotic kinases associated with centromeres and chromosome segregation associate with LANA [24, 55–58]. Thus, these H3 pS10 associated R-loops may provide a unique chromatin environment at the KSHV TR that is both transcriptionally active yet protective for the viral episome during latency. Interesting, DAXX, which has also been shown to colocalize with LANA-NBs, has been implicated in preventing double strand breaks at centromeric R-loops [59]. DAXX may provide both histone chaperone and chromosome protective function along with LANA at the KSHV TRs. DAXX has been shown to suppress R-loop associated DNA-damage at centromeres [59]. Thus, these TR-associated R-loops may provide protective centromere-like function to KSHV genome.

### LANA forms substructures with many of these components

High-resolution microscopic imaging and PLA demonstrate that LANA is in close proximity to many of these epigenetic components, including RNA polymerase, R-loops (S9.6 antibody) and H3pS10 (Figs 4 and 5, Supplemental Figure 3 and Movie 1). The structures formed between LANA and H3pS10 were visualized by high-resolution confocal imaging and 3D reconstructions indicate that these complexes are both large and interwoven, suggesting that they occur over an expansive higher-ordered DNA/chromatin structure. The LANA-NBs have unique epigenetic and morphological features suggesting they form a protective structure important for viral genome transcription, replication and segregation.

### Conclusion

TRC associate R-loops are known to promote DNA recombination and genetic instability, but it is not clear that the R-loops that form at the KSHV TR promote such instability as they appear to occur frequently and viral genetic stability remains intact. While unscheduled R-loops are likely to be threats to genetic integrity, we propose that the R-loops that form at the KSVH TR are part of a programmed mechanism required to maintain the GC-rich repeat structure and promote viral genome integrity. The viral repeat copy number is essential for both viral episome maintenance and tethering to the host metaphase chromosome, as well as for viral lytic cycle DNA cleavage by terminase and virion packaging [60]. We suggest that the R-loops provide a level of complexity to the TR that includes open chromatin with enhancer-like capability as well as protective shell-like properties associated with large nuclear bodies, formed by multiple oligomers of LANA and associated factors, such as DAXX.

## Materials and Methods

### Cells and Drug Treatments

iSLK stable cell lines carrying KSHV Bac16 expressing RFP-LANA or KSHV BAC16-GFP were described previously [27] and cultured in DMEM supplemented with 10% FBS (heat inactivated), 50µg/ml penicillin/streptomycin, 1µg/ml Puromycin, 0.25mg/ml G418, and 1mg/ml Hygromycin B. BCBL1 cells, a stable cell line derived from KSHV^+^ pleural effusion lymphoma (gift from Yan Yuan, UPENN), and BC1 cells (EBV and KSHV positive) were cultured in RPMI-1640 medium supplemented with 10% FBS (heat inactivated) and 50µg/ml penicillin/streptomycin. 293T cells cultured in DMEM with 10% FBS (heat inactivated) and 50µg/ml penicillin/streptomycin. Cells were treated with 0.4µM/DMSO Flavopiridol (Sigma, F3055) or 2µM/DMSO Triptolide (TOCRIS, 3253) dissolved in DMSO for 15 min or 2 hrs as indicated.

### Plasmids and Oligonucleotides

The plasmids containing 2×TR and 8×TR were generous gifts of Dr. Ken Kaye [39]. pCMV3x-FLAG-LANA has been described previously [61]. Oligonucleotides were synthesized by IDT and a complete list is provided in Supplemental Table 1 (Supplemental Table 1).

### ChIP Assay

ChIP assays were performed essentially as previously described [27]. Briefly, cells were collected 1×10^6^ per IP, crosslinked (1% Formaldehyde) with rotation for 15 mins and then quenched by 0.125M glycine for 5 mins. Samples were then lysed in 1 ml SDS lysis buffer (1% SDS, 10 mM EDTA, and 50 mM Tris-HCl, pH 8.0) plus proteinase inhibitor cocktail. Samples were sonicated with a (Diagenode Bioruptor) and diluted 10-fold in IP Dilution Buffer (0.01% SDS, 1.1% Triton X-100, 1.2mM EDTA, 16.7mM Tris pH 8.1, 167mM NaCl, and protease inhibitors cocktail). Appropriate antibody was then added for each sample and rotated overnight at 4°C. Dynabeads (50µl) were added for 2 hours, washed and eluted at 65°C with shaking for 30 mins. Dynabeads were then separated from eluted material by magnet, elute was crosslinked at 65 overnight. DNA was purified using PureLink™ PCR Purification Kit (Invitrogen) according to manufacturer’s instructions. ChIP DNA was assayed by qPCR using primers specific for indicated KSHV regions and quantified as % input.

### DRIP Assay

DRIP assay was performed essentially as described previously [62, 63]. Essentially, 10µg of DNA was incubated with 10ug of S9.6 antibody (Kerafast) and isotype IgG control. Samples were washed with wash buffers, DRIP Buffer (50 mM Tris-HCl, 150 mM NaCl, 5 mM EDTA,1.0% NP-40), DRIP H wash buffer (50 mM Tris-HCl, 500 mM NaCl, 5 mM EDTA,1.0% NP-40, 0.1% Sodium deoxycholate), DRIP Li wash buffer (50 mM Tris-HCl, pH 8.0, 250 mM LiCl, 1 mM EDTA, 0.5% NP-40, 0.5% Na-Deoxycholate) and TE Buffer (100 mM Tris-HCl, 10 mM EDTA 50 mM NaCl) and eluted in elution buffer (50 mM Tris-HCl (pH8.0) 10 mM EDTA 1.0% SDS). DNA was purified using using PureLink™ PCR Purification Kit (Invitrogen) according to manufacturer’s instructions. DRIP DNA was assayed by qPCR using primers specific for indicated KSHV regions and quantified as % input. RNaseH (New England Biolabs, NEB #M0297) treatment was used as per manufacturer’s instructions to validate, as were known cellular positive and negative control primers in 293T cells.

### Indirect immunofluorescence (IF) assay

IF was performed essentially as described previously [17]. Briefly, 1×10^5^ cells seeded at 70% confluence were plated on glass coverslips (treated with Poly-L-Lysine, Corning, 354085) in 24-well plate and incubated overnight in cell culture CO_2_ incubator. Cells were fixed with cold (−20°C) 100% methanol for 10 minutes, washed three times (5 min each wash) with 1xPBS. Fixed cells were incubated 10 min with 0.1M Glycine/PBS and then washed three times with 1xPBS. The cells were permeabilized with 0.3% TritonX-100 (Sigma) in 1xPBS, 10 min. All procedures were performed at room temperature. After washing with PBS, cells were incubated in blocking solution (0.2% fish gelatin, 0.5% BSA in 1xPBS) for 1h. Primary antibodies were diluted in blocking solution and applied on cells for 1h followed with 1xPBS washing. Cells were further incubated with fluorescence-conjugated secondary antibodies AlexaFluor 488 or 594,or 647 (Invitrogen) in blocking solution for 1h, counterstained with 0.5µg/ml DAPI (Sigma) for 5 min, and mounted in ProLong Gold antifade mounting solution (Life Technologies, # P36930). Images were acquired with a Nikon 80i Upright Microscope (Nikon Instruments) using Nikon NIS Elements, AR Advanced Research software, version 6.10.01 (Nikon).

### PLA (Proximity Ligation Assay)

For Proximity ligation assay we followed by the manufactured protocol provided by Sigma: Duolink in Situ Starter Kit mouse/rabbit with reagents (Sigma, Cat. No. DUO92101). The PLA probes from different species, one PLUS (Sigma, DUO9200) and one MINUS (Sigma, DUO92004), matching the host species of primary antibodies. Depended on the conjugated color we used two deferent kits of detection reagents: the Detection reagent Green (495) (Sigma, DUO92014); or Detection reagents Red (594) (Sigma, DUO92008). Briefly, 1×10^5^ cells, 70% confluence, were plated on glass coverslips (treated with Poly-L-Lysine, Corning, 354085) in 24-well plate, for overnight. Cells were fixed with cold (−20°C) 100% Methanol for 10 minutes, washed three times (5 min each wash) with 1xPBS. Fixed cells were incubated 10 min with 0.1M Glycine/PBS and then washed three times with 1xPBS. The cells were permeabilized with 0.3% TritonX-100 (Sigma) in 1xPBS, 10 min. All procedures were performed at room temperature.

After 3 times washing with 1xPBS, cells were incubated in Duolink blocking solution (Sigma, DUO82007) for 1h, at 37°C, followed by Incubation with a pair of primary antibodies (rabbit and mouse), diluted in Duolink Antibody Diluent (Sigma, DUO8208), during 1h at room temperature. After incubation the cells were washed two times, 5 min each, with Duolink Washing buffer-A (Sigma, DUO82046), followed by incubation with PLA probe PLUS and MINUS for 1h, 37°C. Next, the cells were washed 2 times with Buffer-A, followed by Ligation (30 min, 37°C) and Amplification (100min, 37°C), then, the cells were washed with Buffer-B (Sigma, DUO82048), 2 times, 10min each, at room temperature. In the end the cells were mounted in PLA Mounting medium with DAPI (Sigma, DUO2040).

For combination of PLA with IF, after ligation and washing, a secondary AlexaFluor (488) antibodies against the primary anti-LANA antibody was applied. Secondary antibodies were diluted in Antibody diluent, incubated 1h, room temperature, followed by washing with Buffer-A and Amplification step after. Images were acquired with a Nikon 80i Upright Microscope (Nikon Instruments) using Nikon NIS Elements AR Advanced Research software, version 6.10.01 (Nikon).

### Confocal microscopy and image processing

High resolution, confocal images of fixed cells were captured using a Leica TCS SP8 WLL scanning laser confocal microscope and Leica LAS-X software (Leica Microsystems, Inc., Buffalo Grove, IL). Image post-processing included importing into Huygens software for deconvolution (Scientific Volume Imaging, Laapersveld, Hilversum, The Netherlands) followed by 2D maximum projection or 3D reconstruction, iso-surface application and video rendition in LAS-X.

Fixed cell preparations were acquired using a 63X/1.40 oil objective, 6X zoom and a pinhole of 1 AU, with 11–15 z-steps through 3–3.5 µm stacks, resulting in a voxel size of 60 x 60 x 299 nm.

For 3D reconstructions, original confocal files were captured using a Leica TCS SP8 WLL scanning laser confocal microscope (Leica Microsystems, Inc., Buffalo Grove, Ill.). Extended focus Z-stacks of each cell were acquired according to optimized settings based on Nyquist criteria and the resulting files were processed using Huygens deconvolution software (Scientific Volume Imaging, B.V., Hilversum, Netherlands). 3D reconstructions and video animations were then created using the Leica LAS-X 3D software module.

### Centrifugal Elutriation

Centrifugal elutriation using Beckman Coulter AVANTI J-26 XP was used to separate BCBL1 cells into different phases of the cell cycle, with flow rates of 15, 18, 21, 24, 27, 30, 34, 38, 42, and 46 ml/min, as described previously [64].

### Western blot

Briefly, cells (∼1 x 10^6^) were lysed in RIPA buffer (150 mM NaCl, 25 mM Tris, pH 8.0, 1% Na-Deoxycholate, 0.5% SDS, 1% NP-40, 10mM Glycero-phosphate, 1mM Sodium fluoride, 2mM Sodium orthovanadate, 1mM EDTA, and Protease Inhibitors freshly added). An equal amount of protein was resolved using Novex 8-16% Tris-Glycine Gels (Invitrogen) and then transferred onto Nitrocellulose membrane (Millipore) followed with specific antibodies application. Antibody signal was detected using Immobilon Forte Western HRP Substrate (Millipore) and Luminescent Imager 680 (Amersham Bioscience). Primary antibodies were used: anti-HRP-b-Actin (Sigma, A23852), rat anti-LANA HHV8 (Abcam, ab4103), mouse Phospho-RNA Pol II Ser2 (Active Motif, 61083), mouse Phospho-RNA Pol II Ser5 (Invitrogen MA1-460093), rabbit Phospho-HisH3, Ser10 (Invitrogen, 701258).

### Antibodies

The following antibodies were used for immunofluorescence, Co-IP, and Western blotting studies: rat polyclonal anti-HHV8 or anti-LANA [LN53] (Abcam, ab4103), mouse anti-ORF45 (provided by Yan Yuan, UPENN), Rabbit anti-LANA (Novus, NBP3-07279), rabbit Phospho-His H3-Ser10 (Invitrogen, 701258), mouse Phospho-Histone H3 (Ser10) (Invitrogen, MA5-15220), mouse DNA-RNA Hybrid S9.6 (Kerafast, ENH001), mouse Phospho-RNA Pol II Ser2 (Active Motif, 61083), rabbit Phospho-RNA Pol II Ser2 (Abcam, 5095), mouse Phospho-RNA Pol II Ser5 (Invitrogen MA1-460093), rabbit Phospho-RNA Pol II Ser5, (Abcam, 39233), rabbit PCNA (Invitrogen, PA5-27214), Rabbit CTCF (Actif Motif, AB-3216313), RAD21 (Abcam ab217678), Anti-HRP-Flag (Sigma, A8592). The following secondary antibodies were used: AlexaFluor594, or AlexaFluor488, or AlexaFluor647 (Invitrogen), rabbit IgG (Santa Cruz, sc-2027) and mouse anti-Actin-HRP (Sigma, A23852).

### RNA seq analysis

Public data sets from [36] GEO: GSE179727 and from [35] GEO GSE147063 were mapped to the KSVH terminal repeats 2x with a portion of unique region K15 using bowtie 2 [65]with option “--very-sensitive-local” mode so that soft-clipped reads also can be aligned and remain strand-specific.

## Acknowledgments

We thank the Wistar Institute Cancer Center core facilities of Genomics, Bioinformatics, and Imaging. We thank members of the Lieberman lab for their generous assistance, particularly L. MacMullen, S. Soldan, and C. Albitz,. We thank Dr. Ken Kaye for generous gift of p8TR and p2TR.

## Author contributions

Conceptualization: AA, PML

Methodology: AA, AS, OV, JH, JW, MEM, KN, MM

Investigation: AA, AS, OV

Visualization: JH, PML

Supervision: PML

Writing—original draft: AA, PML

Writing—review & editing: AA, PML

## Data and materials availability

All data are available in the main text or the supplementary materials. GSE numbers are provided for the analysis of previously published public data sets. Reagents are available upon request.

---

**Supplementary Fig. S1.**
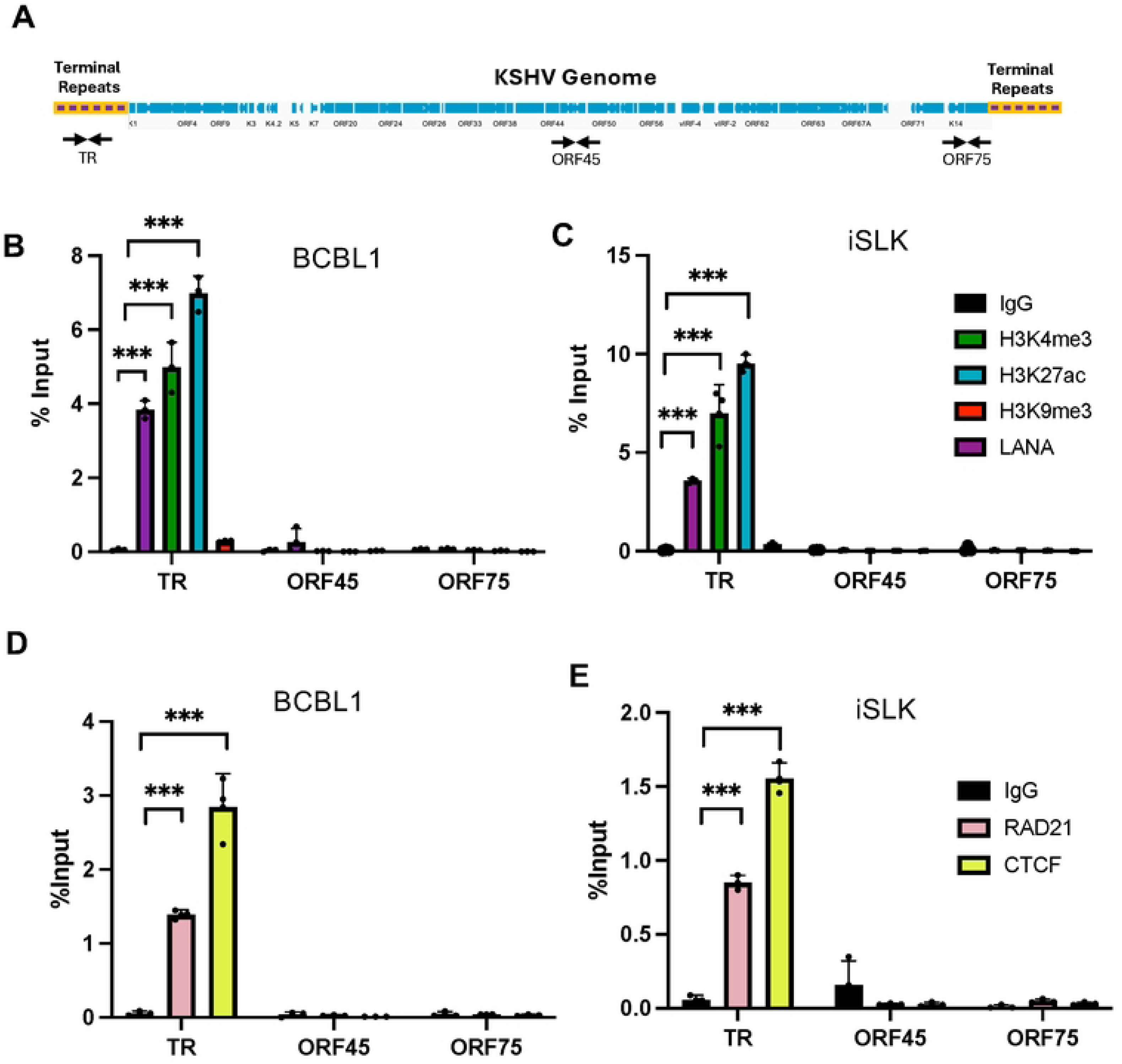
Colocalization of epigenetic marks with LANA at KSHV TR. **A**. Schematic of the KSHV genome showing the terminal repeats (TR) relative to the unique region open reading frames (blue) and primer positions for TR, ORF45, and ORF75. **B.** ChIP-qPCR for histone H3K4me3, H3K27ac, H3K9me3, LANA or control IgG assayed at the TR, ORF45 or ORF75 loci in BCBL1 or iSLK cells. **C.** Same as in panel B, except ChIP antibodies with RAD21, CTCF, or IgG control. ** p<.01, *** p<.001, student 2-tailed t-test, n=3 biological replicates.

**Supplementary Fig. S2.**
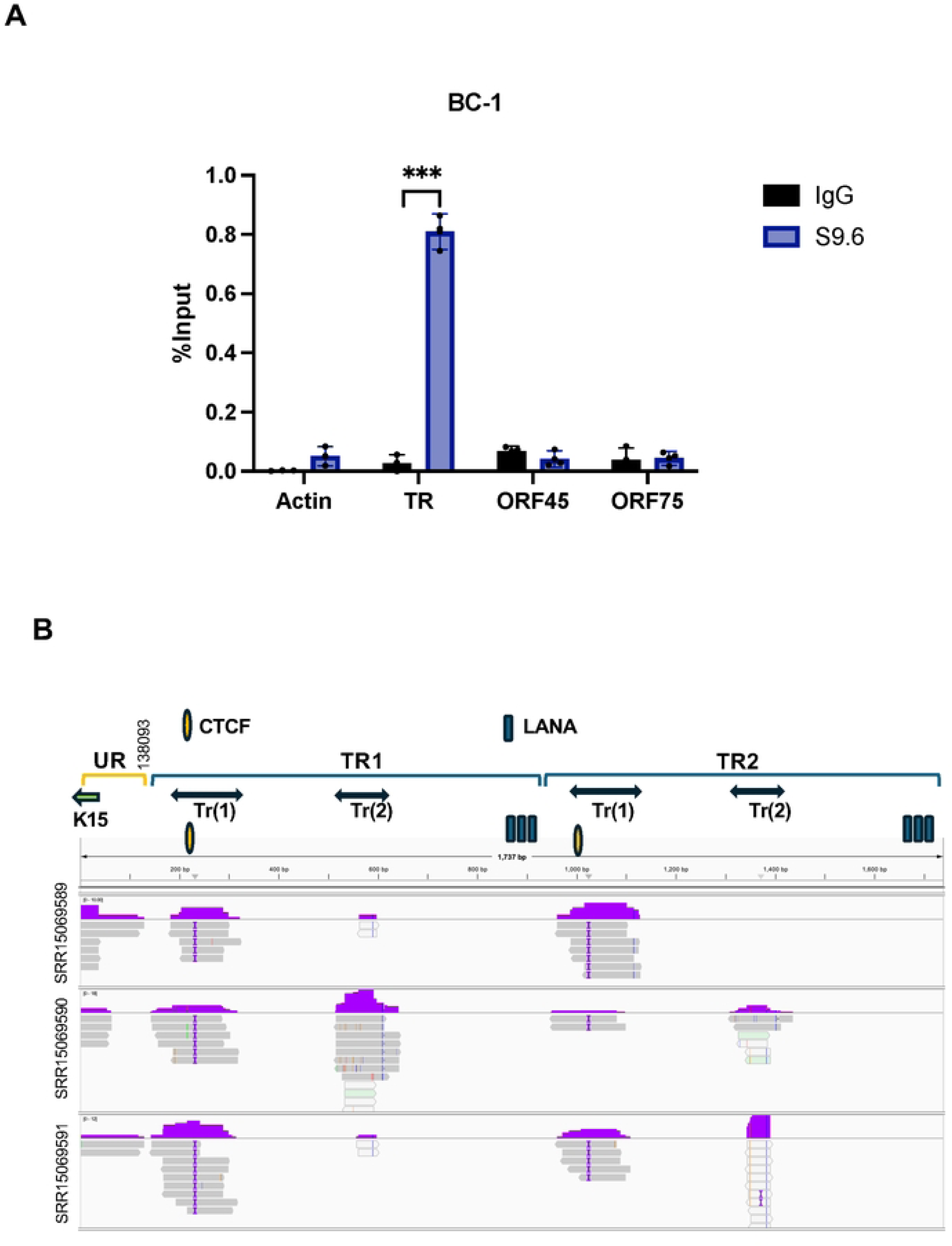
R-loop formation at the KSHV TR. A. DRIP assay with BC1 cells using S9.6 (blue) or control IgG (black) assayed with primers for cellular actin or KSHV TR ORF45 and ORF75. *** p<.001, student two-tailed t-test. B. IGV screen shot of RNA transcripts mapped to KSHV TR region using public data sets for total RNAseq in BCBL1 cells during latent conditions (SRR15069589, SRR15069590, SRR15069591). The reference map consists of a small region of the unique region with K15 and 2 copies of the TR. TR transcripts Tr(1) and Tr(2) are indicated above.

**Supplementary Fig. 3.**
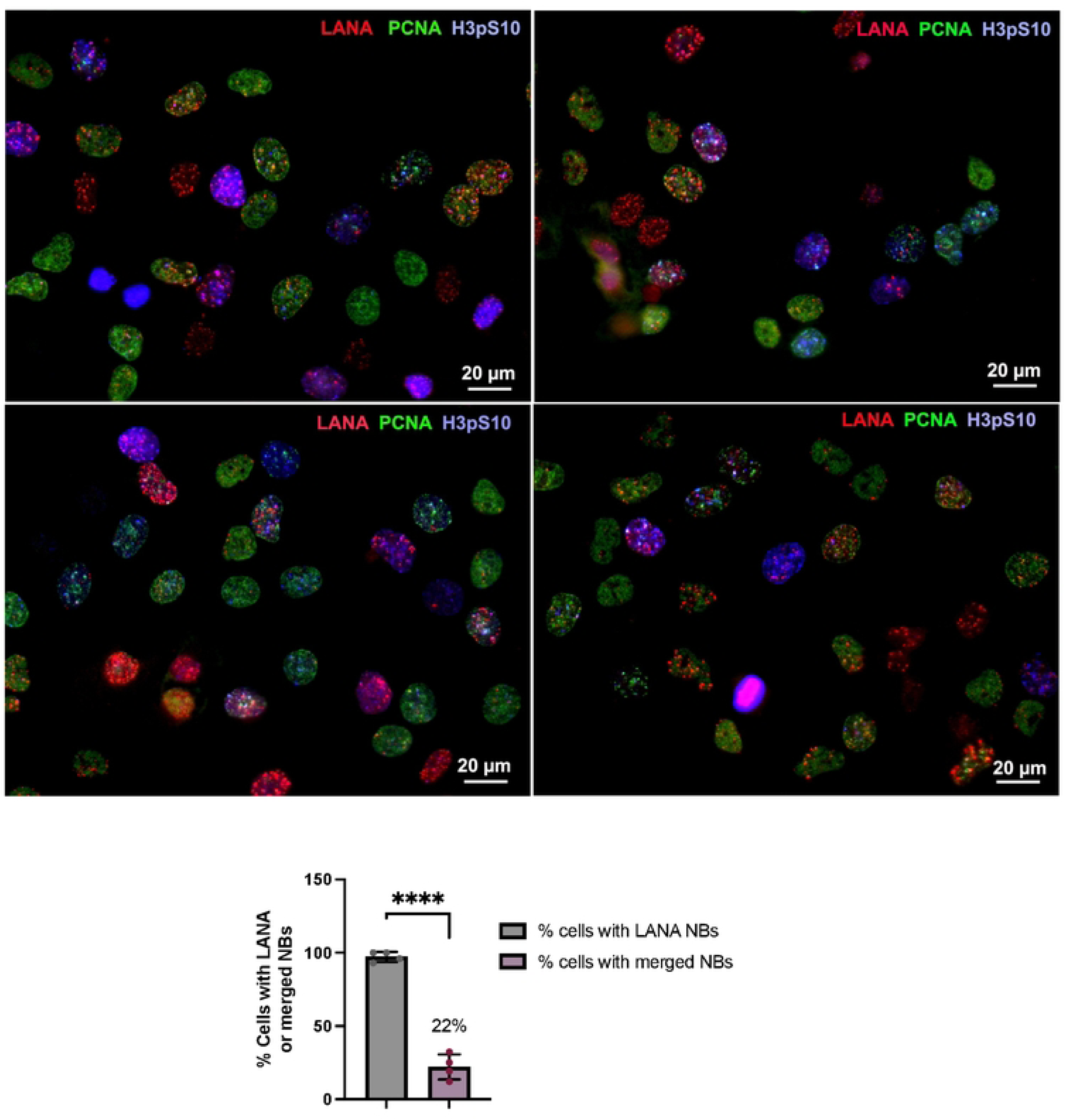
Colocalizations of LANA, H3pS10 and PCNA. Representative images of iSLK cells showing the percentage of cells with LANA and colocalizations with H3pS10 and PCNA. 60x magnification, N=4, total of 126 cells, ****p<.0001, student two-tailed t-test.

**Supplementary Fig. S4.**
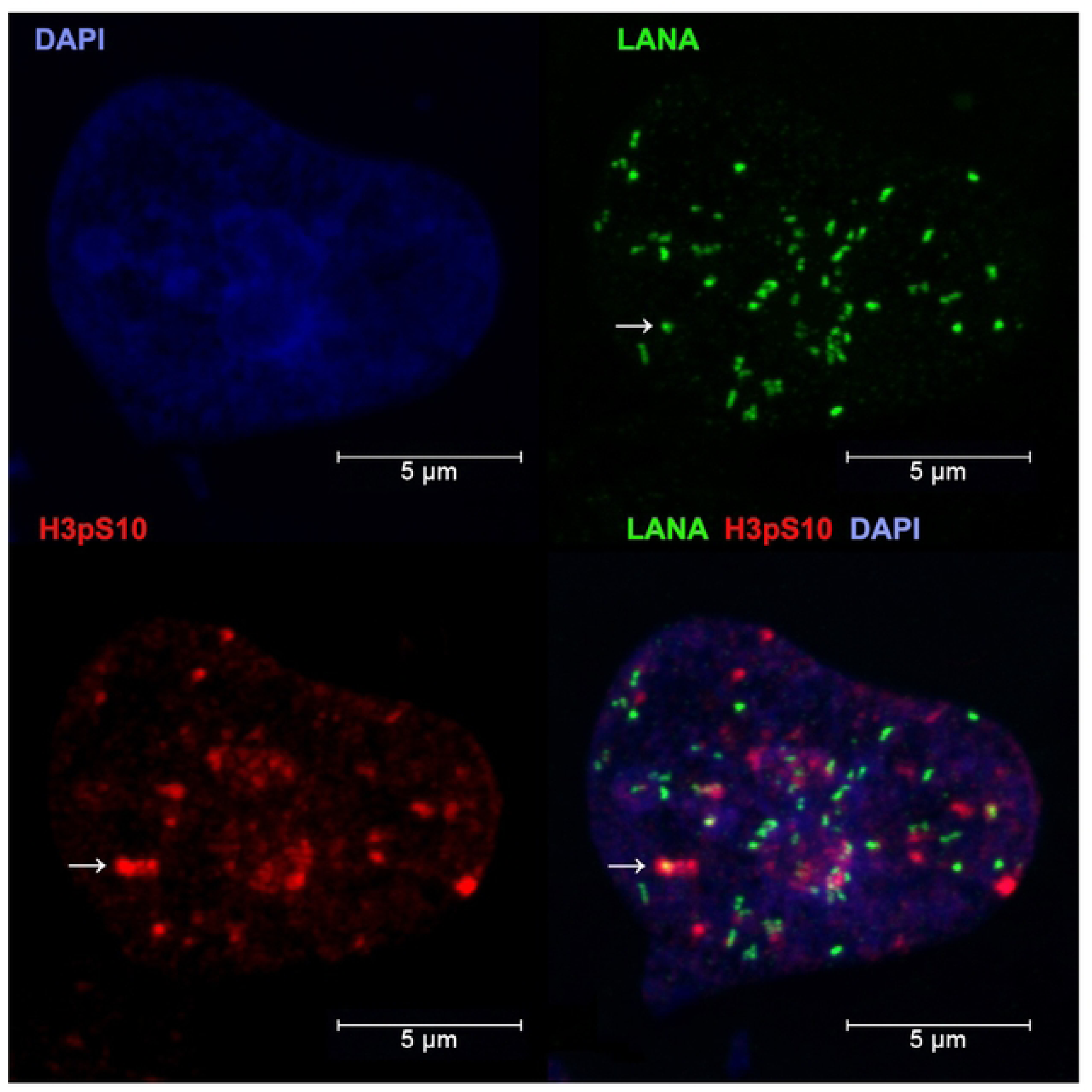
Confocal microscopy IF analysis of H3pS10 and LANA colocalization in iSLK cells. H3pS10 (blue), LANA (red), Dapi (blue).

**Supplementary Fig. S5.**
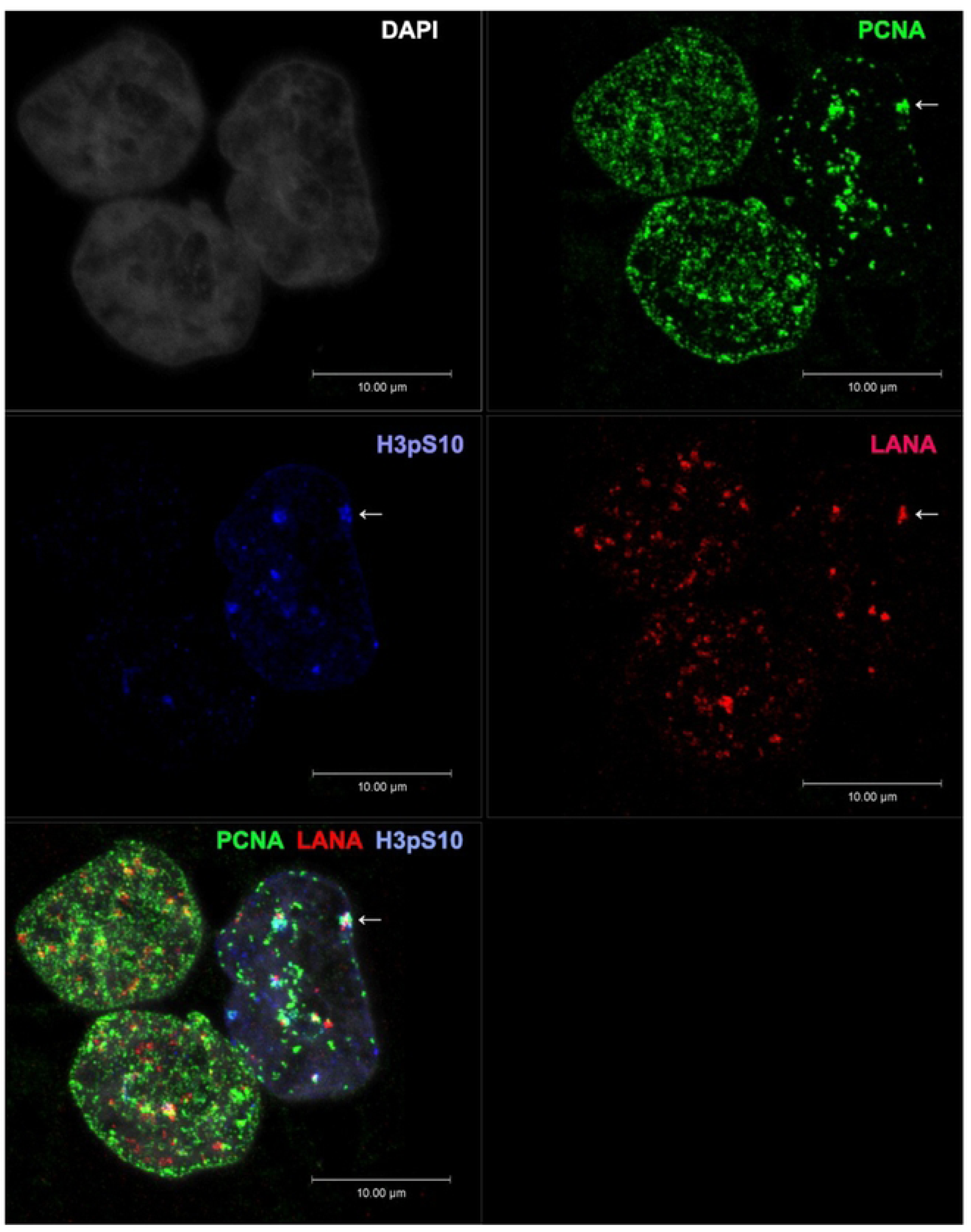
Confocal microscopy IF analysis of PCNA-H3pS10-LANA colocalization in iSLK cells. PCNA (green), H3pS10 (blue), LANA (red).

**Supplementary Fig. 6.**
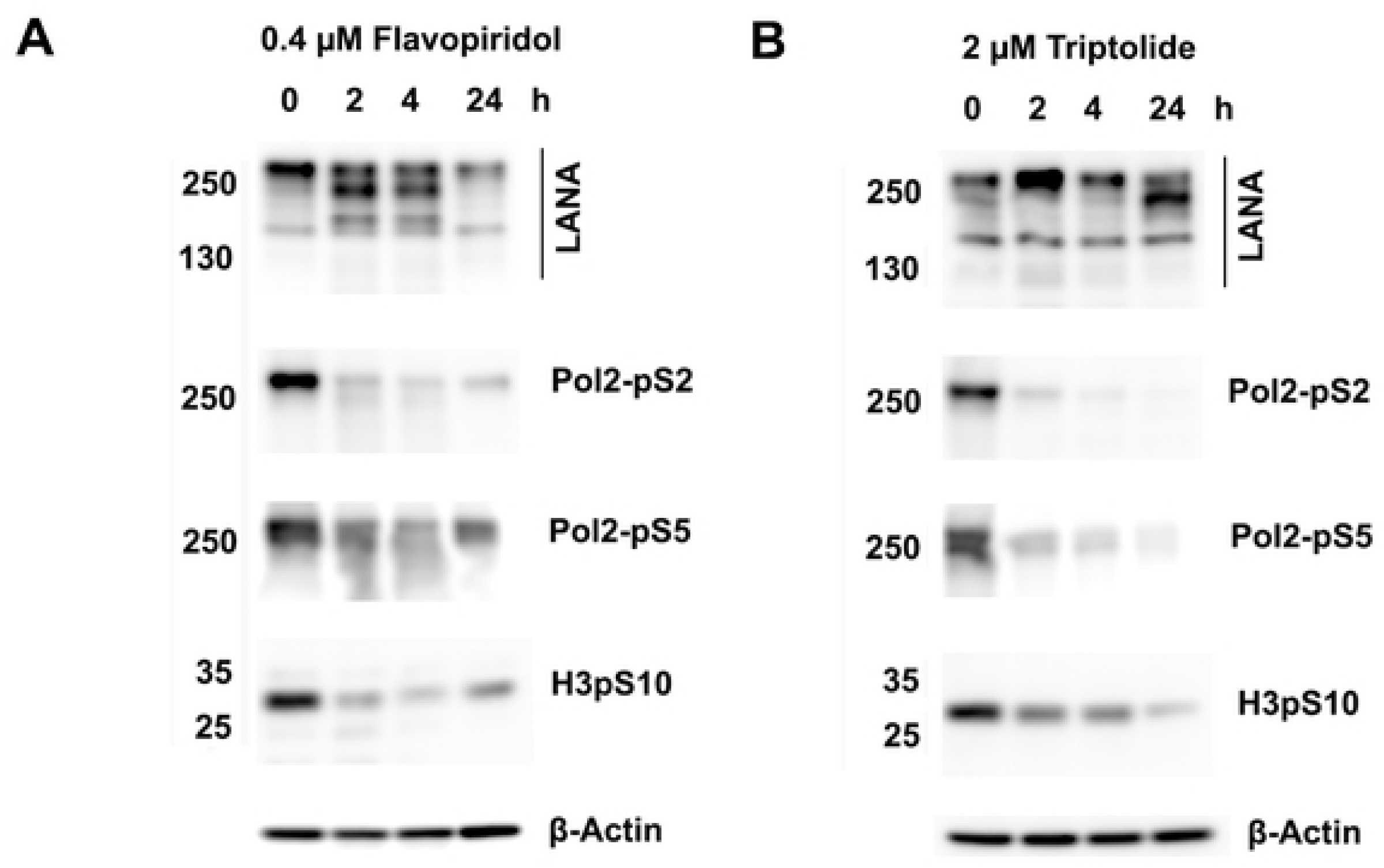
Western blot of BCBL1 cells treated with FVP or triptolide. **A-B.** Western blots of BCBL1 cells treated with 0.4 mM FVP (pane A) or with 2 mM triptolide (panel B) for 0, 2, 4 or 24 hrs and probed for LANA, RNAPII pS2, pS5, H3pS10, or b-actin.

**Supplementary Movie. M1. Confocal microscopy IF analysis of PCNA-H3pS10-LANA colocalization in iSLK cells.** PCNA (green), H3pS10 (blue), LANA (red).

